# Subtly altered topological asymmetry of brain structural covariance networks in autism spectrum disorder across 43 datasets from the ENIGMA consortium

**DOI:** 10.1101/2021.05.05.442735

**Authors:** Zhiqiang Sha, Daan van Rooij, Evdokia Anagnostou, Celso Arango, Guillaume Auzias, Marlene Behrmann, Boris Bernhardt, Sven Bolte, Geraldo F. Busatto, Sara Calderoni, Rosa Calvo, Eileen Daly, Christine Deruelle, Meiyu Duan, Fabio Luis Souza Duran, Sarah Durston, Christine Ecker, Stefan Ehrlich, Damien Fair, Jennifer Fedor, Jacqueline Fitzgerald, Dorothea L. Floris, Barbara Franke, Christine M. Freitag, Louise Gallagher, David C Glahn, Shlomi Haar, Liesbeth Hoekstra, Neda Jahanshad, Maria Jalbrzikowski, Joost Janssen, Joseph A. King, Luisa Lazaro, Beatriz Luna, Jane McGrath, Sarah E. Medland, Ciara Molloy, Filippo Muratori, Declan G.M. Murphy, Janina Neufeld, Kirsten O’Hearn, Bob Oranje, Mara Parellada, Jose C. Pariente, Merel C. Postema, Karl Lundin Remnelius, Alessandra Retico, Pedro Gomes Penteado Rosa, Katya Rubia, Devon Shook, Kristiina Tammimies, Margot J. Taylor, Michela Tosetti, Gregory L. Wallace, Fengfeng Zhou, Paul M. Thompson, Simon E. Fisher, Jan K. Buitelaar, Clyde Francks

## Abstract

Small average differences in the left-right asymmetry of cerebral cortical thickness have been reported in individuals with autism spectrum disorder (ASD) compared to typically developing controls. Although these alterations affect multiple and widespread cortical regional asymmetries, the extent to which specific structural networks might be affected remains unknown. Inter-regional morphological covariance analysis can capture network connectivity relations between different cortical areas at the macroscale level. Here, we used cortical thickness data from 1,455 individuals with ASD and 1,560 controls, across 43 independent datasets of the ENIGMA consortium’s ASD Working Group, to assess hemispheric asymmetries of intra-individual structural covariance networks, using graph theory-based topological metrics. Compared with typical features of small-world architecture in controls, the ASD sample showed significantly altered asymmetry of hemispheric networks involving the fusiform, rostral middle frontal, and medial orbitofrontal cortex, driven by shifts toward higher randomization of the corresponding right-hemispheric networks in ASD. A network involving the superior frontal cortex showed decreased right-hemisphere randomization. Based on comparisons with meta-analyzed functional neuroimaging data, the altered connectivity asymmetry particularly affected networks that subserve working memory, executive functions, language, reading, and sensorimotor processes. Taken together, these findings provide new insights into how altered brain left-right asymmetry in ASD affects specific structural and functional brain networks. Altered asymmetrical brain development in ASD may be partly propagated among spatially distant regions through structural connectivity.

## Introduction

Autism spectrum disorder (ASD) is a childhood-onset condition of neurodevelopmental origin with a prevalence of roughly 1%^1–4^. Individuals with ASD are characterized by social communication and interaction challenges alongside restricted and/or repetitive behaviours causing functional impairment in major areas of life^3^. Language delay is also a common feature of the disorder^5, 6^.

Brain regions important for social cognition and language show lateralized activation in functional neuroimaging studies, in the majority of people^7^. For example, roughly 90% of the adult population has left-hemispheric dominance for word generation tasks, which particularly elicit activation of inferior frontal and temporal cortex^8, 9^, while theory-of-mind tasks typically elicit rightward asymmetrical activation around the temporo-parietal junction^10^. Moreover, increased autism symptom severity has been associated with reduced laterality of activation during language and social cognition tasks^11, 12^. At the structural level, too, studies have reported altered asymmetries of regions of the cortex important for language and/or social cognition, including temporal regions and the fusiform gyrus^13–15^. These findings suggest that altered asymmetrical neurodevelopment may be etiologically linked to ASD behavioral phenotypes.

We recently performed the largest-to-date study of brain structural asymmetry in ASD^13^, analyzing a total of 1,774 affected individuals and 1,809 controls from multiple datasets made available by the ASD working group of the international ENIGMA (Enhancing Neuro-Imaging Genetics through Meta-Analysis) consortium^16, 17^. ASD was most notably associated with widespread alterations of cortical thickness asymmetry, involving the medial frontal, posterior cingulate and inferior temporal cortex. These regions overlapped with those showing altered functional lateralization for language and social cognitive tasks in ASD^11, 12^.

The widespread nature of altered cortical thickness asymmetries in ASD, over multiple non-contiguous regions, raises a new question: do the findings indicate altered asymmetry of topological network organization in ASD? Network organization can be investigated using cortical thickness data from *in vivo*, non-invasive structural MRI, by studying the inter-regional covariance of thickness measures^18–21^. Cortical thickness is a widely-used morphological measure to estimate structural networks^22^, as it relates to underlying features such as the sizes and densities of neurons^23, 24^, as well as functional and white matter connectivity^22, 25^. While it is not fully understood how inter-regional covariation of cortical thickness arises, one prevailing hypothesis is that synapses can have mutually trophic and protective effects on the pre- and post-synaptic neurons involved, such that increased inter-regional connectivity can lead to co-variance at the macro-anatomical level^19^. In addition, synchronous firing between neurons could trigger coordinated synaptogenesis and growth of more highly connected regions^26, 27^.

Neural connections may also propagate pathological processes between spatially distant regions^28^, which has led to a notion of brain disorders as being partly “disconnection syndromes”^29–31^. For example, lower structural covariance based on regional thickness measures from the fronto-temporal cortex has been observed in individuals with ASD relative to typically developing controls, an association which may also be modulated by language and social cognitive abilities^32–35^. However, the regions in these studies were defined by *prior* knowledge of language, whereas alterations of cortical thickness in ASD are more widespread than this^16^. Transcriptome analyses based on postmortem cortical tissue have implicated disrupted biological pathways affecting cell number, cortical patterning and differentiation, axon guidance, synaptic activity and plasticity-related processes in ASD^36, 37^. This also suggests a broader impact on cortical structure beyond core language regions.

Thus far, investigations of altered topological network connectivity in ASD have been impeded by limited sample sizes in relation to subtle effects, and the likely neurobiological heterogeneity of ASD. In addition, studies of altered cross-subject morphological covariance have so far ignored intra-individual structural covariance. The latter reflects structural covariance between different brain regions within each individual (see below)^20, 21, 38^, and can therefore better capture global and regional network characteristics at an individual level.

No previous studies of structural covariance network connectivity in ASD have specifically addressed the possibility of altered network left-right asymmetry at the whole-hemisphere level. Here, we hypothesized that ASD is associated with the reorganization of hemispheric cortical thickness covariance network architecture, such that altered inter-regional connectivity asymmetry could link some of the disparate regions that have previously shown altered asymmetry in separate region-by-region testing^13^. We used structural MRI data from 43 datasets (1,455 ASD patients and 1,560 unaffected controls), collected by members of the ENIGMA consortium’s ASD Working Group, to perform the first graph-based, cortex-wide analysis of structural covariance network asymmetry in ASD. This was followed by functional annotation of affected networks through the use of meta-analyzed functional neuroimaging data, as well as tests relating altered structural network covariance asymmetry within ASD individuals to symptom severity, psychiatric medication use, IQ, age, sex and handedness.

## Material and Methods

### Datasets and participants

Structural T1-weighted brain MRI-derived data were available via the ENIGMA-ASD Working Group ^16^. After data quality control (see below), there were 1,455 individuals with ASD (mean age: 15.65 years, range 2-64 years, 1,213 males) and 1,560 healthy controls (mean age: 16.09 years, range 2-64 years, 1,179 males) across 43 separate datasets (**Table 1**). Datasets were collected as separate studies between 1994 and 2013, when DSM-IV and DSM-IV-TR were the common classification systems; clinical diagnosis of ASD was made according to DSM-IV criteria. For each dataset, all subjects were diagnosed by a clinically experienced and board certified physician/psychiatrist/psychologist. Data on DSM-IV subtypes of ASD were not collated by the ENIGMA ASD working group. For each of the data sets, all relevant ethical regulations were complied with, and appropriate informed consent was obtained for all individuals.

**Table 1.** Characteristics of the 43 datasets of the ENIGMA Autism Spectrum Disorder working group that were used in this study.

| Dataset no. | Dataset name | N total | N cases (M/F) | N controls (M/F) | Mean age (min, max) | Scanner type | Field strength |
| --- | --- | --- | --- | --- | --- | --- | --- |
| 1 | ABIDE_CALTECH | 31 | 13/1 | 13/4 | 29.05 (17.5,56.2) | Siemens Trio | 3T |
| 2 | ABIDE_LEUVEN_2 | 35 | 12/3 | 15/5 | 14.16 (12.1,16.9) | Philips Interna | 3T |
| 3 | ABIDE_MAX_MUN | 51 | 21/3 | 24/3 | 26.47 (7,58) | Siemens Verio | 3T |
| 4 | ABIDE_NYU | 184 | 68/10 | 81/25 | 15.29 (6.47,39.1) | Siemens Allegra | 3T |
| 5 | ABIDE_OLIN | 36 | 17/3 | 14/2 | 16.81 (10,24) | Siemens Allegra | 3T |
| 6 | ABIDE_PITT | 58 | 26/5 | 23/4 | 19.17 (9.33,35.2) | Siemens Allegra | 3T |
| 7 | ABIDE_SBL | 30 | 15/0 | 15/0 | 34.37 (20,64) | Philips Interna | 3T |
| 8 | ABIDE_SDSU | 37 | 14/1 | 16/6 | 15.04 (8.67,37.7) | GE MR750 | 3T |
| 9 | ABIDE_STANFORD | 40 | 16/4 | 16/4 | 9.96 (7.53,12.94) | GR Signa | 3T |
| 10 | ABIDE_TCD | 55 | 24/1 | 30/0 | 16.66 (9.3,25.91) | Philips Achieva | 3T |
| 11 | ABIDE_UM_1 | 126 | 48/14 | 41/23 | 12.82 (8.07,20.9) | GE Signa | 3T |
| 12 | ABIDE_UM_2 | 31 | 14/1 | 15/1 | 15.27 (11.14,26.8) | GE Signa | 3T |
| 13 | ABIDE_USM | 100 | 59/0 | 41/0 | 21.34 (8.2,50.22) | Siemens Trio | 3T |
| 14 | ABIDE_YALE | 55 | 20/8 | 19/8 | 12.72 (7,17.83) | Siemens Magnetom | 3T |
| 15 | ABIDEII-BNI | 57 | 28/0 | 29/0 | 38.11 (18,64) | Philips Ingenia | 3T |
| 16 | ABIDEII-EMC | 41 | 18/2 | 19/2 | 8.32 (6.4,10.66) | GE MR750 | 3T |
| 17 | ABIDEII-ETH | 31 | 11/0 | 20/0 | 22.93 (13.83,30.67) | Philips Achieva | 3T |
| 18 | ABIDEII-GU | 98 | 39/8 | 26/25 | 10.71 (8.06,13.91) | Siemens TriTim | 3T |
| 19 | ABIDEII-IP | 52 | 14/7 | 9/22 | 20.66 (6.05,46.6) | Siemens TriTim | 1.5T |
| 20 | ABIDEII-IU | 39 | 15/4 | 15/5 | 24.38 (17,54) | Philips Achieva | 3T |
| 21 | ABIDEII-KKI | 199 | 35/15 | 94/55 | 10.34 (8.01,12.99) | Philips Achieva | 3T |
| 22 | ABIDEII-NYU_1 | 72 | 38/4 | 28/2 | 10.05 (5.22,34.76) | Siemens Allegra | 3T |
| 23 | ABIDEII-OHSU | 92 | 29/7 | 27/29 | 10.98 (7,15) | Siemens Skyra | 3T |
| 24 | ABIDEII-OILH | 39 | 12/1 | 17/9 | 23.54 (18,31) | Siemens TriTim | 3T |
| 25 | ABIDEII-SDSU | 57 | 26/7 | 22/2 | 12.99 (7.4,18) | GE MR750 | 3T |
| 26 | ABIDEII-TCD | 41 | 19/0 | 22/0 | 15.38 (10,20) | Philips Achieva | 3T |
| 27 | ABIDEII-USM | 32 | 15/2 | 12/3 | 21.36 (9.12,38.86) | Siemens TriTim | 3T |
| 28 | BRC | 44 | 17/0 | 27/0 | 14.75 (10,18) | GE Signa HDx | 3T |
| 29 | Barcelona | 52 | 29/2 | 20/1 | 11.97 (7.25,17.08) | Siemens Trio | 3T |
| 30 | Dresden | 45 | 18/3 | 20/4 | 35.33 (21.1,56.8) | Siemens Trio | 3T |
| 31 | FAIR | 81 | 33/6 | 27/15 | 11.47 (7.84,15.94) | Siemens Magnetom | 3T |
| 32 | FSM | 80 | 20/20 | 20/20 | 4.1 (1.83,6) | GE Signa | 1.5T |
| 33 | MRC | 137 | 67/0 | 70/0 | 26.96 (18,45) | GE Signa HDx | 3T |
| 34 | PITT_1 | 56 | 11/3 | 34/8 | 16.32 (8,36) | Siemens Allegra | 3T |
| 35 | PITT_2 | 89 | 38/6 | 39/6 | 16.97 (8,36) | Siemens Allegra | 3T |
| 36 | ParelladaHGGM | 66 | 33/2 | 30/1 | 12.52 (7,18) | Philips Intera | 1.5T |
| 37 | TCD_2 | 27 | 10/0 | 17/0 | 16.94 (12.7,24.8) | Philips Achieva | 3T |
| 38 | TORONTO_1 | 177 | 70/20 | 45/42 | 11.83 (3.3,20.8) | Siemens Trio | 3T |
| 39 | TORONTO_2 | 192 | 99/41 | 28/24 | 11.03 (2.48,21.67) | Siemens Trio | 3T |
| 40 | UMCU_1 | 57 | 25/3 | 27/2 | 14.3 (7.11,24.72) | Philips | 1.5T |
| 41 | NIJMEGEN2 | 68 | 27/18 | 15/8 | 26.29 (18,40) | Siemens Avanto | 1.5T |
| 42 | NIJMEGEN3 | 92 | 36/4 | 43/9 | 9.54 (6.07,12.27) | Siemens Avanto | 1.5T |
| 43 | NIJMEGEN1 | 33 | 14/3 | 14/2 | 14.95 (12.28,18.02) | Siemens Trio | 3T |

Total scores from the Autism Diagnostic Observation Schedule-Generic (ADOS)^39^ were available for 704 individuals with ASD. The presence or absence of comorbid conditions had been recorded for 200 individuals with ASD, and there were 43 affected individuals diagnosed with at least one other co-occurring condition (e.g., attention deficit hyperactivity disorder, obsessive-compulsive disorder, depression, anxiety and/or Tourette’s syndrome^16^). Data on the presence or absence of medication use at the time of scanning (i.e., current use of psychiatric treatment drugs prescribed for ASD or comorbid Binary psychiatric conditions) were available for 612 individuals with ASD, of whom 173 were current users. Data on IQ were available for 1,210 of the ASD individuals. Cases from the entire ASD spectrum were included, but only 61 cases had IQ<70 (cases: mean IQ=103.49, SD=19.74, min=34, max=149). categorical data on handedness were available for 599 ASD individuals (551 right-handed, 48 left-handed).

There were different assessment and recruitment processes for controls across the datasets, but the overwhelming majority were typically developing at the time of scanning, and no controls met criteria for a diagnosis of ASD. Only 18 controls had IQ<70. In these individuals the exclusion of an ASD diagnosis was performed by a senior child psychiatrist/physician. All eighteen of these were from the FSM data set and were clinically diagnosed with idiopathic intellectual disability. Amongst all 1,303 controls with IQ data, the mean IQ was 111.75, SD=14.73, min=31, max=149.

### Image acquisition and processing

Structural T1-weighted brain MRI scans were collected at each separate study site, using a variety of different scanners and protocols at field strengths of either 1.5 or 3 Tesla (**Table 1**). Following this heterogeneous image acquisition, all sites applied the same harmonized protocol from the ENIGMA consortium (http://enigma.ini.usc.edu/protocols/imaging-protocols) for data processing and quality control^16, 17^. FreeSurfer^40^ (version 5.3) was used to derive mean cortical thickness measures for each of 68 cortical regions (34 per hemisphere) defined by the Desikan-Killiany atlas^41^. The default ‘recon-all’ pipeline of FreeSurfer was used, which incorporates renormalization^40^. Parcellations of cortical regions were visually inspected following the standardized ENIGMA quality control protocol (http://enigma.ini.usc.edu/protocols/imaging-protocols). Briefly, web pages were generated with snapshots from internal slices, as well as external views of the segmentations from different angles. For subcortical structures, the protocol also consisted of visually checking individual images, plotted from a set of internal slices. Values derived from incorrectly labelled structures were excluded. Furthermore, any data points exceeding 1.5 times the interquartile range, as defined per site and diagnostic group, were visually inspected, and any errors resulted in excluded values.

Specifically for the present study, we also excluded any individuals with missing thickness data for at least one cortical region, as the analyses required all regions for comparability of networks across individuals. We also excluded datasets with fewer than 15 controls, as variation within the control group of each dataset is important for calculating intra-individual covariance (see below). These steps resulted in 43 datasets being included in the present study, with the sample numbers given above.

### Construction of intra-individual hemispheric structural covariance networks

Within each dataset, regional cortical thickness values were used to separately construct left-hemispheric and right-hemispheric structural covariance networks for each individual (**Fig. 1**). Such networks are comprised of nodes and edges, in which each cortical region represents a node. The edge between each pair of regions in a given individual was calculated with respect to the standard deviations for those regional measures calculated from control individuals^20, 21^. The formula was as follows^20, 21^:

**Figure 1.**
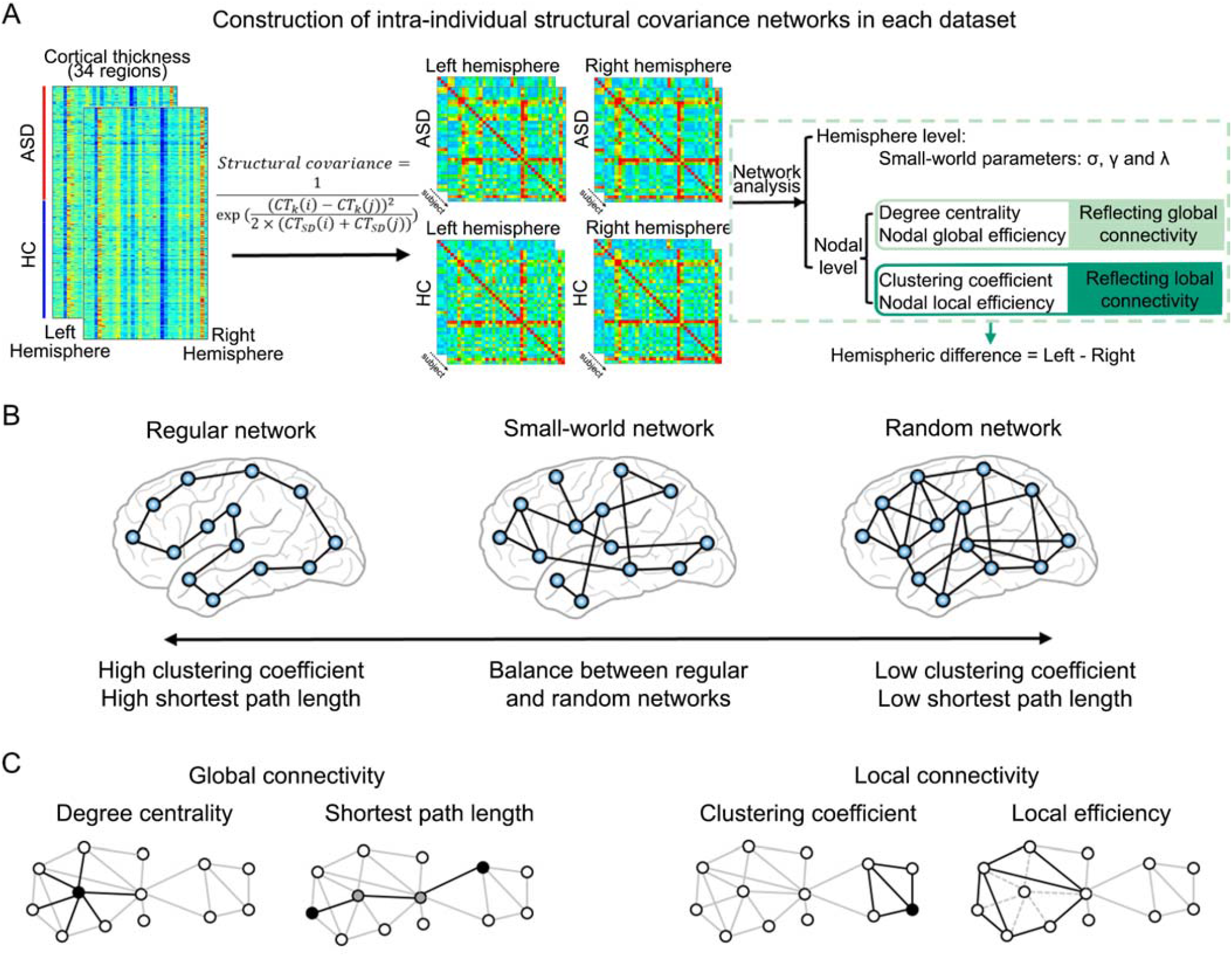
Schematic workflow of this study. (A) Flowchart of the procedure used in the current study. We first constructed intra-individual, intra-hemispheric structural covariance networks in each dataset using regional cortical thickness data. Then, for each individual, we computed graph theory metrics at the global and nodal levels using the intra-hemispheric networks. Finally, we calculated individual-level hemispheric differences for each metric, to examine case-control differences of topological network asymmetry. (B) Small-world network model. At the whole-hemisphere level, we estimated network integration and segregation using small-world parameters. A regular network is characterized by a high clustering coefficient and long shortest path length, corresponding to high local specialization and low global integration. In contrast, a random network has a low clustering coefficient and short shortest path length, corresponding to low local specialization and greater global integration. A small-world model reflects a balance between the extremes of local specialization versus global integration. (C) At the nodal level, we examined four graph theory measures: degree centrality and nodal global efficiency both measure global connectivity from/to a given node, whereas the cluster coefficient and nodal local efficiency reflect local connectivity from/to that node. Abbreviations: ASD: autism spectrum disorder; HC: healthy control; SD: standard deviation.

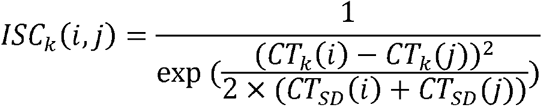

where *ISC*_*k*_ (*i,j*) represents the intra-individual structural covariance between region *i* and region *j* in individual *k. CT*_*k*_ (*i*) and *CT*_*k*_ (*j*) represent the cortical thicknesses of regions *i* and *j* in individual *k. CT*_*SD*_ (*i*) represents the standard deviation of cortical thickness in region *i* across all control individuals in a given dataset.

For each individual, this approach yielded two separate 34×34 matrices, one for the left hemisphere and one for the right hemisphere, each representing a network of intra-hemispheric structural connectivity, with 561 edges in each network. Edges were then binarized using a sparsity threshold S=0.4, i.e., only the top 40% of strongest edges (i.e., 224 edges) within each separate network were given the value 1, and the remainder 0, in order to reduce spurious connectivity. The sparsity threshold S=0.4 retained network connectedness, such that at least 88% of nodes (30 nodes) remained connected with at least one other node in all left- and right-hemisphere networks in all individuals (i.e., a maximum number of four isolated nodes in any network). Small-world organization was also retained (minimum small-worldness scalar σ=1.0001 in any network, see below). This approach ensured that all left- and right-hemisphere networks, in all individuals, had the same number of nodes (34) and edges (224), and enabled us to perform subsequent analyses with reference only to relatively high-level, reliable connectivity^42^.

### Hemisphere-level network properties

We used the Brain Connectivity Toolbox^43^ and GRETNA^44^ toolbox to calculate topological network indices (**Fig. 1**). At the whole-hemisphere level, we used small-world parameters to measure the balance between network integration and segregation. Small-worldness can be quantified by the clustering coefficient and shortest path length^45^. Based on these measures, a network can be described as regular, random or small-world. A regular network is characterized by a high clustering coefficient and high shortest path length, indicating high local specialization (high local efficiency) and low global integration (low global efficiency). In contrast, a random network has a low clustering coefficient and low shortest path length, corresponding to low local specialization (low local efficiency) and high global integration (high global efficiency). In general, human brain networks are organized in an optimized, small-world fashion^18, 46^, with an intermediate balance between regular and random properties, i.e., a large number of short-range connections coexist with a smaller number of long-range connections.

The clustering coefficiet *c_i_* of the node *i* was defined as^45^

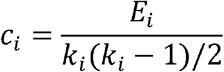

where *E_i_* represents the number of existing edges among the neighbors of node *i*. *k_i_* denotes the actual number of neighbors of node *i*, thus the denominator quantifies the number of all possible edges among the neighboring nodes. The clustering coefficient *C* of a whole hemispheric network was then defined as the mean clustering coefficient across all nodes in that network, separately per individual and hemisphere.

The shortest path length *l_i_* of a node *i* was defined as^45^

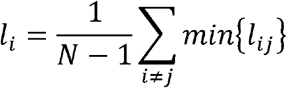

where min{*l*_*ij*_} represents the shortest path length between node *i* and *j. N* represents the number of nodes in the network. The shortest path length *L* of a network was then defined as the mean shortest path length between any pair of nodes in the network, calculated separately per individual and hemisphere.

Next, we generated 100 random networks with the same number of nodes (34), edges (224), and degree distribution as the real networks^47^ to calculate a normalized clustering coefficient γ=*C*/*C*_rand_ and normalized shortest path length λ =*L*/*L*_rand_ ^45^, in which *C*_rand_ and *L* _rand_ were defined as the mean clustering coefficient and mean shortest path length across randomly generated networks. The small-world index σ was calculated as σ=γ/λ, which should be greater than 1 in small-world networks, and whose minimum value was 1.0001 across the networks of all individuals and hemispheres (see above)^45^.

These procedures resulted in three hemisphere-level connectivity metrics for each individual and hemispheric network: the normalized clustering coefficient γ the normalized shortest path length λ and the small-world index σ

### Node-level network properties

For each of the 34 nodes, separately per individual and hemisphere, we calculated four measures: the degree centrality and nodal global efficiency (both indicate connectivity globally from/to a given node) and the clustering coefficient and nodal local efficiency (both indicate local connectivity from/to a given node) ^42, 48, 49^.

Specifically, the degree centrality of node *i* was defined as the sum of all existing edges between that node and all other nodes in the network, reflecting the importance of that node in network information communication.

The global efficiency of node *i* (*E*_*glob*_) indexes information transfer from itself to all other nodes in the entire network^48^, computed as the reciprocal of the shortest path length *l_i_* :

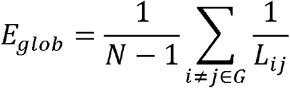

where *L*_*ij*_ is the shortest path length between node *i* and node *j* in network *G. N* is the number of nodes in the network.

The clustering coefficient measures the extent of local density of connections for a given node (see the formula further above, where node-level clustering coefficients were calculated as a step towards the hemisphere-level clustering coefficient).

The local efficiency of node *i* (*E*_*loc*_) corresponds to the efficiency of information flow within the local environment^48^, which is defined as

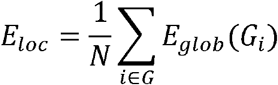

where *G_i_* represents the subgraph composed of the nearest neighbors of node *i*.

### Hemispheric asymmetry

To quantify the asymmetry of each separate network metric within each individual, we calculated the hemispheric difference (HD):

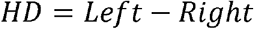

Therefore, a positive value of HD represents a leftward asymmetry for a given metric, while a negative HD represents a rightward asymmetry for that metric. (Note that the widely-used asymmetry index (Left-Right)/(Left+Right) would be less well suited to the present study, as Left and/or Right could sometimes take the value zero for the metrics defined above, in which case this index would take extreme or undefined values.)^50^

### Statistical analysis

We used linear mixed effects random-intercept models (‘fitlme’ function in MATLAB version 2016a (The Mathworks Inc.) to test for case-control differences, across all datasets simultaneously, but separately for each network metric HD. All models included the same fixed effects, i.e., diagnosis (case versus control status), age and sex, plus a random effect indicating which of the 43 datasets an individual was from, as shown in the following formula:

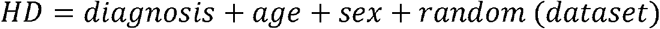

The random effect ‘dataset’ adjusted for all variables that differed between datasets, including scanner type and field strength (Table 1). The *t* values derived from the “diagnosis” factor were used to compute Cohen’s *d* effect sizes^51^

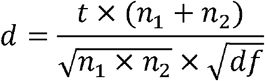

where *n*_1_ and *n*_2_ are the numbers of cases and controls, and *df* represents the degrees of freedom. *df* = *obs* − (*x*_1_ + *x*_2_), where *obs* equals the number of observations *x*_1_, the number of groups and *x*_2_the number of factors in the model.

Permutations (N=10,000) were used to test the significance of each case-control diagnosis effect on each HD metric, by randomly assigning the diagnosis labels across individuals, separately within each dataset while maintaining the same numbers of cases and controls within each dataset, prior to mega-analysis across datasets. Shuffling was carried out within datasets separately because intra-individual covariance was calculated with reference to the variance in matched controls (see above). The empirical p value was obtained for each diagnosis effect on each HD metric by counting the number of unsigned *t* values in the permutation analysis that were greater than the unsigned *t* value for the data with the real case-control labels, and dividing that number by the total number of permutations (N=10,000). For the 3 network-level metric HDs, significance was determined by the *p* values for diagnosis effects with Bonferroni correction of 0.05/3. For the node-level network metric HDs, significance was determined for diagnosis effects using false discovery rate (FDR) correction for 34 nodes, with threshold pFDR<0.05/4 (due to testing four nodal-level network measures).

### Directions of topological network asymmetry changes

For any HDs showing significant case-control differences in the main analysis above, we used linear mixed effects models to examine separately the corresponding left and right metrics to understand the unilateral effects. Models with the same fixed and random effects were used as above, and again *t* values and Cohen’s *d* effect sizes for the diagnosis term were extracted/derived from each of the models. Empirical *p* values were calculated based on 10,000 permutations, as above. As this was a post hoc analysis to further describe any specific alterations of asymmetry in cases, we did not perform multiple testing correction for these analyses.

### Associations with ASD severity, medication, IQ, age, sex or handedness

For each topological network HD that showed significant associations with case-control status in the main analysis, we used separate linear mixed effects models to examine possible relationships between these HDs and ASD severity, medication use, IQ, age, sex or handedness within the ASD individuals only. These analyses would inform whether major aspects of case heterogeneity were related to the relevant topological network HDs.

Autism severity was based on the total ADOS scores of ASD individuals (N=704):

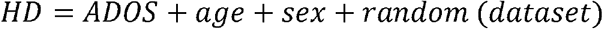

The presence/absence of current psychiatric medication use was coded as a binary predictor variable (0=no medication, 2=medication) (N=612)

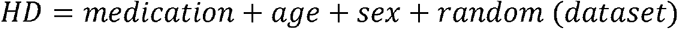

IQ was tested for association with HDs within the 1,210 ASD individuals with IQ data. IQ was coded as a continuous predictor variable:

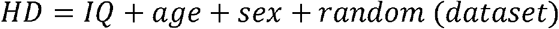

Age and sex were tested for associations with HDs within 1,455 ASD individuals (age as a continuous variable, sex as a binary variable:

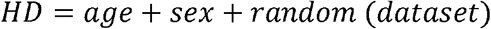

Handedness was tested for association with HDs within 599 ASD individuals that had data on this trait. Handedness was coded as a binary predictor variable (1=right handedness, 2=left handedness).

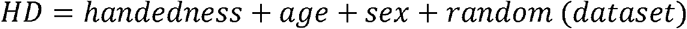

For each of these analyses, permutations (N=10,000) were used to test the significance of each effect of interest, by shuffling the relevant variable (either ADOS scores, medication use, IQ, age, sex or handedness) across case individuals, separately within each dataset, prior to mega-analysis across datasets. *P* values were obtained by counting the number of unsigned *t* values in the permutation tests that were greater than the actual unsigned *t* values for the real data, and dividing by the total number of permutations (N=10,000) for each separate analysis. *P* values were FDR-corrected at 0.05 for multiple testing over the number of network HDs that showed significant effects in the main analysis (i.e., seven network HDs, see Results). We did not additionally correct for multiple testing over the six variables of interest (ADOS scores, medication use, IQ, age, sex or handedness) as this was an exploratory analysis to describe how each case heterogeneity variable might impact the network HDs.

### Descriptive edge-level analysis

For each specific node that showed a significant case-control difference of degree centrality asymmetry in the main analysis, we extracted the intra-hemispheric structural connectivity values (i.e., one value for each edge) linking this ‘seed’ node to all the 33 other nodes, separately from each hemispheric structural covariance network of each individual (this time without thresholding and binarization for sparsity, see above). For each matched pair of left and right edges, we then calculated the HD (again as Left-Right). The same linear mixed effects random-intercept model as the main analysis was used to examine each edge HD as the dependent measure across individuals, and 10,000 permutations were again used to assess the empirical two-tailed significance of the effect of diagnosis. Separately for each relevant node, the *p* value was FDR-corrected at 0.05 for multiple testing over the 33 edges connecting to that node.

### Cognitive functional annotation based on Neurosynth

To indicate the potential cognitive functions of regions that showed altered degree centrality asymmetry, we used the online platform Neurosynth^52^ (https://neurosynth.org/) which includes meta-analytic brain maps based on input data from >14,000 human functional neuroimaging studies. As of February 2021 there were 1,307 maps in the database, representing different terms that capture diverse cognitive functions. Each map indicates a pattern of brain activation linked to a given term, through semantically-related words that occurred in the papers describing those studies. The large size of the database has been shown to compensate for any imperfect assignment of activations to particular cognitive domains or tasks^52^. This approach therefore provides a data-driven alternative to assigning brain regional functions by *ad hoc*, selective citations of limited numbers of papers from the literature.

Separately for each cortical region with significantly altered asymmetry of degree centrality in the node-level analysis, plus all regions linked to them by edges that showed significant alterations of asymmetry in ASD in the edge-level analysis (see above), we mapped these regions to bilateral masks in MNI standard space. The resultant binary masks were then used as input to identify region-associated cognitive terms through the Neurosynth “decoder” function. Finally, cognitive terms with correlations >0.2 were visualized on a word-cloud plot, with sizes scaled according to their correlations with the corresponding meta-analytic maps generated by Neurosynth, while excluding anatomical terms, non-specific terms (e.g. ‘Tasks’), and one from each pair of virtually duplicated terms (such as ‘Words’ and ‘Word’).

### Sensitivity analyses

To assess robustness with respect to the sparsity threshold 0.4 that was used in the main analysis, we repeated the analyses under varying sparsity thresholds ranging from 0.25 to 0.5 (with an interval of 0.01). At the lowest threshold, a minimum 79% of nodes (27 nodes) were connected to at least one other node in all hemispheric networks in all individuals (maximum seven unconnected nodes out of 34). We then computed the area under the curve for each network metric HD over the range of sparsity thresholds. The area under the curve for a given network HD *Y*, calculated over the sparsity threshold range of S_1_ (0.25) to S_n_ (0.5) with interval of Δ S (0.01), was computed as

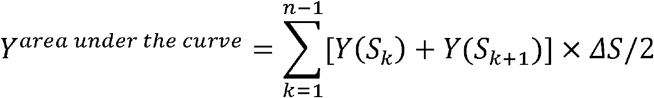

Case-control differences in the area under the curve for each network metric HD were then tested separately using the same mixed effects random-intercept model as the main analysis:

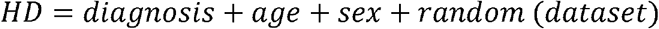

To assess whether non-linear age could have an impact on case-control differences of network HDs, we added a non-linear age term ‘zage^2^’, i.e., (age-mean_age)^2^ as a fixed effect in the linear mixed effects model to test each association:

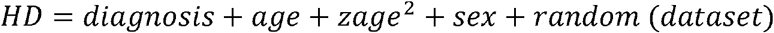

Permutations (N=10,000) were used as in the main analysis, to determine the empirical significance of case-control differences for each network HD metric.

## Results

### Hemisphere-level network asymmetries

None of the three hemisphere-level metric HDs (i.e., the normalized clustering coefficient γ, the small-world index σ, or the normalized shortest path length λ) showed significant differences between individuals with ASD and controls (all p>0.05). A non-significant trend effect of diagnosis was observed for a leftward shift in λ asymmetry in ASD (Cohen’s *d*=0.06, p=0.10; **Supplementary Table 1**). Unilateral analysis of each hemisphere showed that ASD was nominally associated with reduced λ in the right hemisphere (Cohen’s *d*=−0.07, unadjusted p=0.04), but not in the left hemisphere (Cohen’s *d*=0.004, p=0.92), which hints at a more efficient global information transmission and a shift towards randomization of right hemisphere networks in ASD (**Supplementary Table 2**).

### Node-level network measures

We mapped the Cohen’s *d* effect sizes of associations between node-level network measure HDs and ASD over the whole cerebral cortex (**Fig. 2**). Effect sizes were low, ranging from −0.15 (nodal global efficiency HD of fusiform) to 0.14 (degree centrality HD of superior frontal cortex) (**Supplementary Tables 3-6**). Among node-level metric HDs, the degree centrality asymmetries of three regions, namely fusiform (Cohen’s *d*=−0.14, p<0.0001), rostral middle frontal cortex (Cohen’s *d*=−0.13, p=0.0007) and superior frontal cortex (Cohen’s *d*=0.14, p=0.0003), were significantly associated with ASD after FDR correction (**Fig. 2 and Supplementary Table 3**). In addition, nodal global efficiency HDs of four regions, namely fusiform (Cohen’s *d*=−0.15, p=0.0001), rostral middle frontal cortex (Cohen’s *d*=−0.13, p=0.0001), superior frontal cortex (Cohen’s *d*=0.13, p=0.0007) and medial orbitofrontal cortex (Cohen’s *d*=−0.11, p=0.001), were significantly associated with ASD after multiple testing correction (**Fig. 2 and Supplementary Table 4**). Overall, reduced leftward lateralization was observed in network measure HDs of the fusiform, rostral middle frontal and medial orbitofrontal cortex in ASD (**Supplementary Tables 3 and 4**). Superior frontal cortex showed reduced rightward asymmetry of both degree centrality and global efficiency HDs in ASD (**Supplementary Tables 3 and 4**). There were no significant associations between ASD and the HDs of the nodal clustering coefficient or nodal local efficiency after FDR correction (**Fig. 2 and Supplementary Tables 5 and 6**).

**Figure 2.**
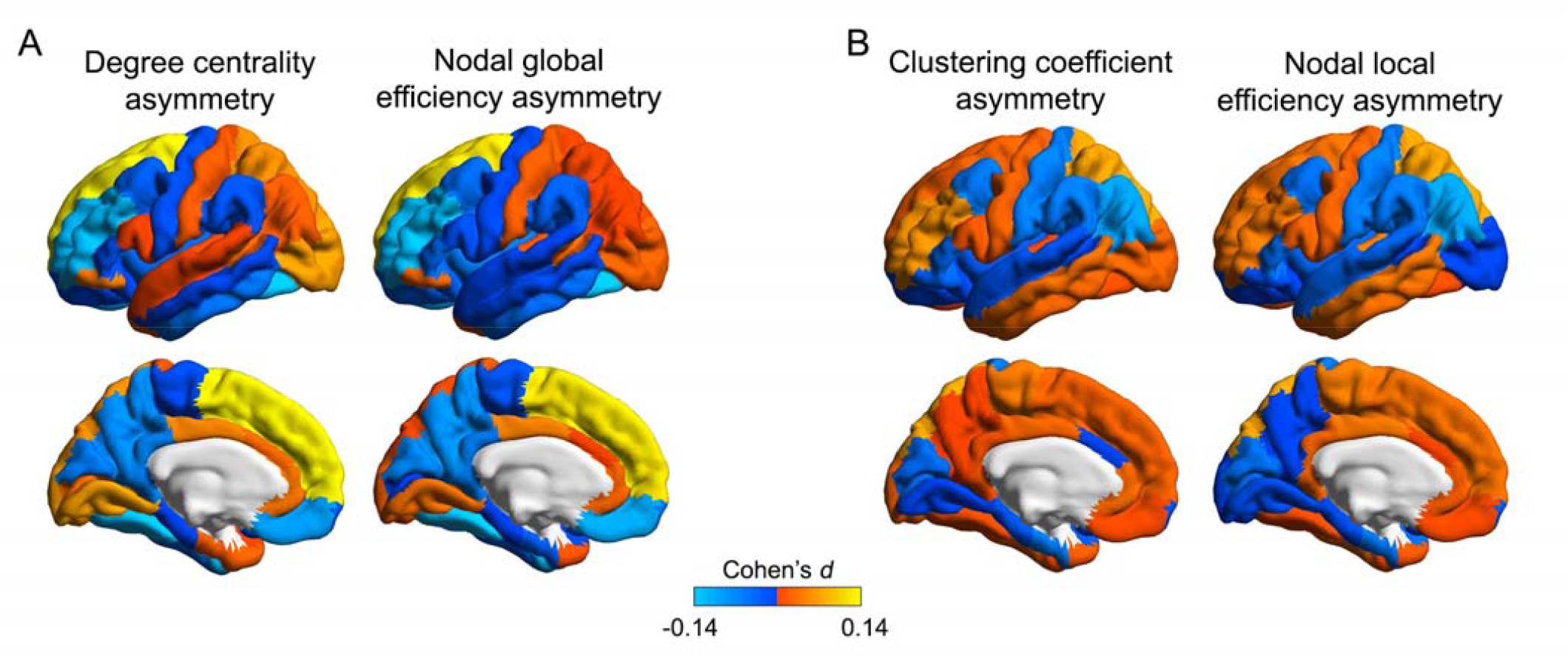
Cohen’s *d* effect sizes of ASD case-control associations for node-level topological asymmetries. (A) Effect sizes from ASD case-control analysis of node-level topological metric asymmetries that reflect global connectivity of each node, i.e., degree centrality and nodal global efficiency. (B) Effect sizes from ASD case-control analysis of nodal-level topological metric asymmetries that reflect local connectivity of each node, i.e., the clustering coefficient and nodal local efficiency. Positive effect sizes (orange-yellow) indicate shifts towards greater leftward or reduced rightward asymmetry in ASD compared to controls, and negative effect sizes (blue) represent shifts towards greater rightward asymmetry or reduced leftward asymmetry in ASD compared to controls.

Further investigating the significant effects on asymmetry in unilateral analyses, the effects on degree centrality asymmetries of the fusiform and rostral middle frontal cortex, and nodal global efficiency asymmetries of the fusiform and medial orbitofrontal cortex, involved right-sided increases, thus resulting in reduced leftward topological asymmetries (**Supplementary Table 7**). For the effect on nodal global efficiency asymmetry of the rostral middle frontal cortex, bilateral increases were observed in ASD, but more so in the right than left hemisphere, consistent with reduced leftward lateralization in ASD individuals relative to controls. The effects on degree centrality and nodal global efficiency asymmetries of the superior frontal cortex involved bilateral decreases in ASD, but more so in the right hemisphere, consistent with reduced rightward asymmetry of these metrics in ASD (**Supplementary Table 7**).

### Clinical severity, medication, IQ, age, sex and handedness

For the 7 network HDs that showed significant case-control differences in our main analysis, we found no significant associations with autism symptom severity (total ADOS scores) (*p*s>0.05; **Supplementary Table 8**). There also were no significant associations of current medication use with these metric HDs after FDR correction (**Supplementary Table 9**). Medication status showed a nominally significant association with the degree centrality HD of the fusiform (Cohen’s *d*=−0.22, unadjusted p=0.04), and a marginal trend with fusiform nodal global efficiency HD (Cohen’s *d*=−0.19, unadjusted p=0.06). There were no significant associations of IQ with the network HDs within ASD individuals (*p*s>0.05; **Supplementary Table 10**). Age showed a significant positive association with the nodal global efficiency HD of the medial orbitofrontal cortex (t=2.36, unadjusted p=0.006; **Supplementary Table 11**). There were no significant associations between network HDs and sex (**Supplementary Table 12**) or handedness (**Supplementary Table 13**).

### Descriptive edge-level analysis

The degree centrality of each node provides a metric of its hemisphere-wide connectivity. For the three regions that showed significant associations between their degree centrality HDs and ASD in the main analysis, i.e., fusiform, rostral middle frontal and superior frontal cortex, we performed descriptive edge-level analysis of case-control associations. Four edges linked to the fusiform cortex showed significant associations with ASD after FDR correction, which linked to the rostral middle frontal (Cohen’s *d*=−0.12, p=0.0004), cuneus (Cohen’s *d*=−0.14, p=0.0005), medial orbitofrontal (Cohen’s *d*=−0.11, p=0.002), and postcentral regions (Cohen’s *d*=−0.13, p=0.0006; **Fig. 3A and Supplementary Table 14**). These edges all showed reduced leftward asymmetry in ASD relative to controls (**Supplementary Table 14**). A significant association was also observed between ASD and connectivity asymmetry between the rostral middle frontal and three other regions, which were the inferior parietal region (Cohen’s *d*=−0.13, p=0.0004), fusiform (Cohen’s *d*=−0.12, p=0.0004), and precuneus (Cohen’s *d*=−0.17, p<0.0001) after FDR correction (**Fig. 3B and Supplementary Table 15**). All of these effects involved lower leftward asymmetry in ASD compared to controls. In addition, connectivity between the superior frontal and paracentral cortex showed a significant association with ASD (Cohen’s *d*=0.12, p=0.001; **Fig. 3C and Supplementary Table 16**). This connectivity showed lower rightward asymmetry in ASD compared to controls. In total, among the nine regions with altered connectivity asymmetry in ASD according to edge-level analysis, four were among those associated with altered cortical thickness asymmetry as previously found in separate region-by-region testing in the ENIGMA ASD data^13^.

**Figure 3.**
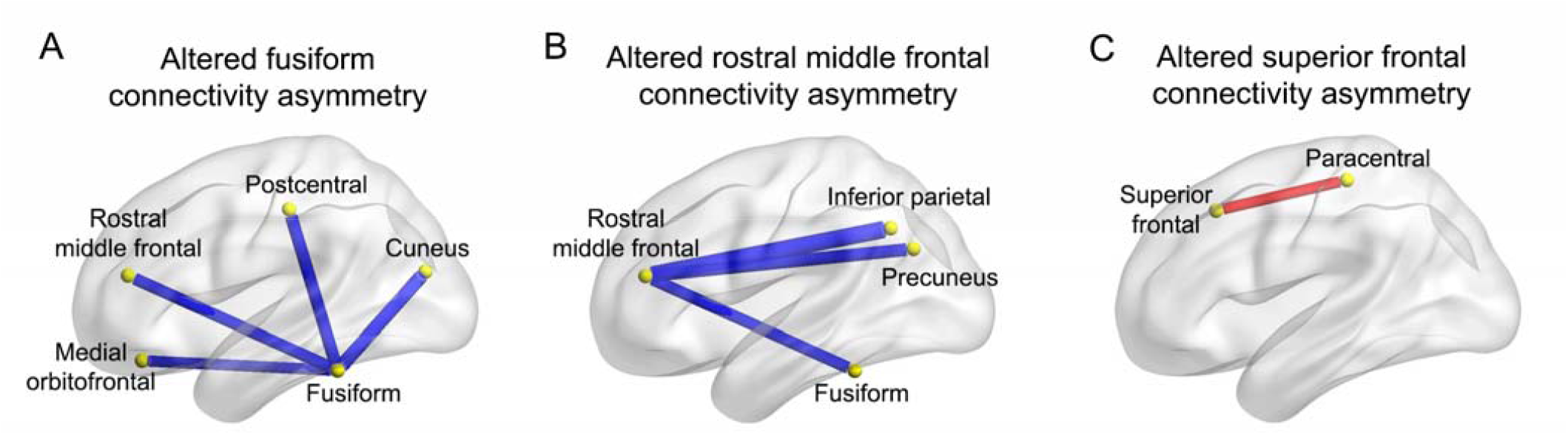
Altered asymmetry of connectivity linking to the nodes with significant alterations of degree centrality asymmetry in ASD. (A) Altered asymmetry of connectivity linked to the fusiform in ASD. (B) Altered asymmetry of connectivity linked to the rostral middle frontal cortex in ASD. (C) Altered asymmetry of connectivity linked to the superior frontal cortex in ASD. The yellow nodes indicate the brain regions. Red indicates a significant edge-level, reduced rightward asymmetry of connectivity in ASD compared to controls, and blue indicates an edge-level, reduced leftward asymmetry of connectivity in ASD compared to controls.

### Functional annotation of networks with altered lateralized connectivity in ASD

The most prominently shared functional annotation for all three networks that showed associations of degree centrality asymmetry with ASD was ‘working memory’ (**Fig. 4 and Supplementary Table 17**). However, each network also had additional cognitive annotations. Disrupted asymmetry of fusiform connectivity involved cortical regions that are especially active during executive control, reading and motor tasks (**Fig. 4A and Supplementary Table 17**). Regions with altered connectivity asymmetry linked to the rostral middle frontal cortex were associated with executive, reading and attention tasks (**Fig. 4B and Supplementary Table 17**). Finally, alteration of superior frontal connectivity asymmetry involved regions associated with executive and sensorimotor tasks (**Fig. 4C and Supplementary Table 17**).

**Figure 4.**
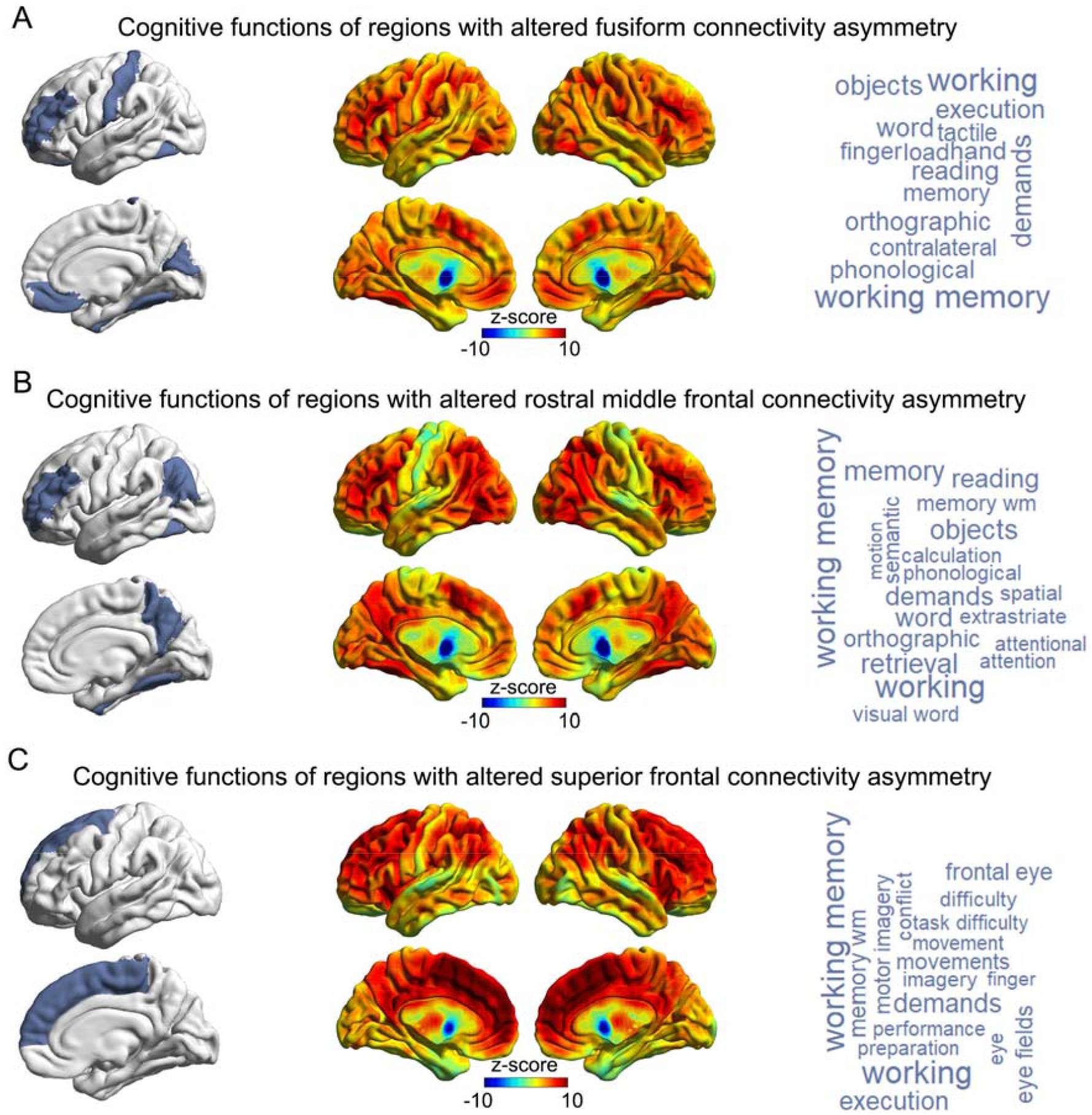
Cognitive functions associated with cortical regions showing altered connectivity asymmetry. Meta-analyzed fMRI data were used to functionally annotate cortical regions showing altered connectivity asymmetry with the fusiform (A), rostral middle frontal (B) or superior frontal (C) cortex. Left panels indicate the regions showing alterations of lateralized connectivity, which were used as input masks to the decoder function of Neurosynth (see Methods). Middle panels show the brain co-activation maps corresponding to the input masks. Right panels show the cognitive terms corresponding to the co-activation maps, in word-cloud plots. The font sizes of the cognitive terms indicate their map-wide correlations with the co-activation maps (correlation coefficients are in Supplementary Table 17).

### Sensitivity analyses

Across the defined range of sparsity thresholds (0.25-0.5), all of the associations remained significant between ASD and the asymmetries of degree centrality and nodal global efficiency, for the regions of fusiform, rostral middle frontal, superior frontal and medial orbitofrontal cortex (**Supplementary Table 18**); this indicated that the findings were robust to threshold selection.

After adding a non-linear age^2^ term to the linear mixed effects models, the effects of case-control status on the 7 affected network HDs remained significant and largely unaffected (**Supplementary Table 19**).

## Discussion

We used a cortex-wide, graph-based approach to investigate differences of topological network asymmetries between individuals with ASD and unaffected controls across 43 datasets of the international ENIGMA consortium’s ASD Working Group. We found significantly lower asymmetry of topological network measures which may reflect altered information transfer in individuals with ASD relative to controls, specifically involving the fusiform, prefrontal and orbitofrontal cortex. The findings were largely driven by a shift to greater randomization of right hemispheric network organization in ASD. The affected structural covariance networks included prefrontal, parietal, posterior cingulate and paracentral cortical regions. Data-driven functional annotation, using meta-analyzed fMRI data, consistently identified working memory as a function that may be especially affected by network asymmetry alterations in ASD, consistent with executive function challenges characteristic of ASD^53–55^.

The network-level findings provide a new understanding of the widespread, dispersed topography of altered cortical thickness asymmetry in ASD that was reported in a previous ENIGMA-ASD study (which used separate region-by-region testing^13^). In other words, some of the altered regions can now be understood in terms of their embedding within specific structural networks that show altered asymmetry in ASD, and have particular functional annotations (more on this below). In general, many cognitive processes involve a degree of left-right hemispheric dominance in the healthy brain^7^, so that the typical asymmetry pattern in the population is likely to be an optimal form of brain organization. It follows that alterations of network-level asymmetry may have functional consequences. As ASD is a childhood-onset disorder, and the majority of individuals in this study were children, the present findings provide further evidence that altered lateralized neurodevelopment is subtly disrupted in ASD. The network-level findings further imply that inter-regional connections may propagate disrupted cortical thickness asymmetries among sets of spatially distant cortical regions, i.e., intra-hemispheric topographic connectivity may help to shape spatial patterns of cortical pathology in ASD.

The effect sizes in this study were small, with Cohen’s *d* ranging from −0.15 to 0.14. These findings indicate that case-control group average differences in structural network asymmetry are very subtle in ASD, and of similar magnitude to those reported in previous ENIGMA consortium studies of brain regional anatomy and asymmetry in ASD^13, 16^. Future studies may apply normative modelling^56^ or clustering^57^ approaches to identify subgroups of individuals with highly atypical structural network asymmetry, and these may constitute etiological subgroups of ASD. MRI-based regional cortical thickness measures are fairly crude biological readouts, affected by numerous possible underlying factors, including the degree of myelination^58^, as well as the numbers and densities of different types of cells and dendritic processes^59–61^. Therefore it also remains possible that subtly altered network asymmetry at the macro scale in ASD reflects more substantial alterations at finer levels of analysis. For example, neurite orientation dispersion and density imaging has been used to study grey matter microstructural asymmetries *in vivo*^62^, or the ratio of T1w and T2w images in grey matter can be used to create an estimate of cortical myelin content^63^. Future *post mortem* studies of cortical histology and gene expression may also reveal microstructural and/or molecular alterations, but there is currently limited data available from homotopic regions of the two hemispheres, as many brain banks assign the left and right hemispheres into distinct storage and analysis protocols^64^.

Three specific cortical regions had node-level degree centrality asymmetries that were significantly altered in ASD: the fusiform, rostral middle frontal, and superior frontal cortices. Our meta-analyzed fMRI-based annotations implicated a range of functions mapping to each of the affected networks involving these regions (**Fig. 4**), which included working memory and other executive function-related annotations in common across the networks, but also language-related, reading-related, and sensorimotor annotations. Language delay is a common feature of ASD^5, 6^, and the disorder is also associated with reduced left-hemisphere language dominance^11^. Numerous reports based on behavioral, neurophysiological, neuroimaging or histopathological data have also reported atypical motor system development in ASD^65^. Our findings may therefore indicate that alterations of specific right-hemisphere structural networks underlie some of the language- and motor-related deficits in ASD. These functional annotations, that were based on meta-analyzed fMRI data from other cohorts, motivate future studies of brain-behaviour correlations using neuroimaging and behaviour data from the same affected individuals.

The fusiform gyrus is especially known to show right-lateralized activation in response to face-related perception^66–69^, which is important in social interactions. Reduced rightward functional asymmetry for face processing has been associated with ASD^70^, so face processing may be one aspect of cognition that is disrupted by increased randomization of a right-hemispheric structural network that includes the fusiform gyrus. The rostral middle frontal cortex (dorsolateral prefrontal cortex), has been proposed to act as a coordinating hub in cognitive control tasks, as part of a frontal-parietal network^71^. This region has been shown to be abnormally active in the left hemisphere in ASD relative to typically developing controls in a recent meta-analysis of cognitive control tasks^72^. Resting-state fMRI data have also suggested a rightward shift in asymmetry of executive control networks in ASD^73^. Moreover, white matter network analysis has suggested that individuals with ASD exhibit a greater age-related increase in global efficiency involving the right dorsolateral prefrontal cortex than typically developing controls^74^. The superior frontal cortex is known as a core region of the default mode network, which can show altered functional asymmetry in ASD^73^. Abnormal lateralization of functional connectivity between the superior frontal gyrus and temporal cortex has also been reported in ASD, and associated with language and social deficits^12^. Our findings further support altered lateralization of superior frontal cortex connectivity in ASD, demonstrated here on a structural level.

For the medial orbitofrontal cortex, there was a significant association of its nodal global efficiency asymmetry with ASD, but no significant association with its degree centrality asymmetry, and we therefore did not include it in fMRI-based annotation and edge-level analysis. The medial orbitofrontal cortex was the only cortical region to show altered asymmetry of both cortical thickness and surface area in individuals with ASD, in a previous ENIGMA consortium study that tested it separately region-by-region (not in a network context)^13^. Another study found that alterations in structural covariance between inferior frontal cortex and the left orbitofrontal cortex was modulated by language ability within ASD individuals^33^, suggesting a possible contribution of the Orbitofrontal cortex to communication deficits in ASD.

Within ASD individuals, we found no associations of the affected network asymmetry metrics with autism symptom severity, psychiatric medication usage, IQ, sex or handedness. Age showed an association with one network metric HD, i.e., the nodal global efficiency HD of the medial orbitofrontal cortex, but apart from this single effect, we were unable to link structural network asymmetries to the within-case phenotypic variables available in the current study. Deeper phenotyping may be needed to understand the relevance of structural connectivity asymmetry alterations in terms of clinical heterogeneity^75, 76^. For example, only total ADOS scores were available through the consortium (rather than subscores that reflect different behavioural dimensions)^39^, and data on medication usage and comorbidities were limited to relatively small subsets of the overall data (see Methods). Future longitudinal studies may help to characterize atypical developmental trajectories of asymmetry patterns in ASD, and capture causal and dynamic processes of structural asymmetry alterations over the course of the disorder. It is also possible that altered structural connectivity will not map onto any identifiable symptom domains of ASD, but rather reflects a shared susceptibility mechanism across various individuals with heterogeneous presentations of ASD, and potentially other diagnoses too.

In conclusion, this consortium study identified small group-average differences between ASD individuals and unaffected controls in specific aspects of the asymmetry of hemispheric structural connectivity networks. The affected networks mapped most consistently to working memory as a function that is influenced by alterations of network connectivity asymmetry in ASD. These findings help to elucidate altered cortical thickness asymmetry in ASD in terms of hemispheric network architecture, and suggest that neurodevelopmental alterations of brain asymmetry in ASD may propagate via structural connectivity.

## Acknowledgements

We thank the participants of all studies who have contributed data to the ENIGMA-ASD working group (http://enigma.ini.usc.edu/ongoing/enigma-asd-working-group/)^16^. This research was funded by the Max Planck Society (Germany). This study was further supported by the ENIGMA Center for Worldwide Medicine, Imaging & Genomics grant (NIH U54 EB020403) to Paul Thompson, and further supported by the Innovative Medicines Initiative Joint Undertaking under grant agreement number 115300 (EU-AIMS) and 777394 (AIMS-2-TRIALS), resources of which are composed of financial contribution from the European Union’s Seventh Framework Programme and Horizon2020 programs and the European Federation of Pharmaceutical Industries and Associations (EFPIA) companies’ in-kind contribution. The Canadian samples were collected as part of the POND network funded by the Ontario Brain Institute (Anagnostou/Lerch). Boris Bernhardt acknowledges research support from the National Science and Engineering Research Council of Canada (NSERC Discovery-1304413), the Canadian Institutes of Health Research (CIHR FDN-154298, CIHR PJT-174995), SickKids Foundation (NI17-039), Azrieli Center for Autism Research (ACAR-TACC), BrainCanada, FRQ-S, and the Tier-2 Canada Research Chairs program. Neda Jahanshad is partially funded by NIH R01MH117601, P41EB015922, R01AG058854, R01MH121246, R01MH111671 and is MPI of a research grant from Biogen Inc, for work unrelated to the contents of this manuscript.

## Contributions

Z.S. and C.F. conceived the study. E.A., C.A., G.A., M.B., G.B.F., S.C., R.C., E.D., C.D., F.L.S.D., S.D., C.E., S.E., D.F., J.F., J.F., D.L.F., C.M.F., L.G., S.H., L.H., N.J., M.J., J.J., J.A.K., L.L., B.L., J.M., F.M., D.G.M.M., K.O., B.O., M.P., A.R., P.R., K.R., D.S., M.J.T., M.T., G.L.W., and F.Z. recruited, assessed and scanned the subjects, and performed MRI processing and quality control. D.v.R. organized the database. Z.S. performed the statistical analysis. Z.S. and C.F. wrote the manuscript. All authors contributed edits and approved the content of the manuscript. C.F. directed the study.

## References

1. Baird G, Simonoff E, Pickles A, Chandler S, Loucas T, Meldrum D et al. Prevalence of disorders of the autism spectrum in a population cohort of children in South Thames: the Special Needs and Autism Project (SNAP). Lancet 2006; 368(9531):210–215.

2. Christensen DL, Baio J, Van Naarden Braun K, Bilder D, Charles J, Constantino JN et al. Prevalence and Characteristics of Autism Spectrum Disorder Among Children Aged 8 Years--Autism and Developmental Disabilities Monitoring Network, 11 Sites, United States, 2012. MMWR Surveill Summ 2016; 65(3):1–23.

3. Diagnostic and statistical manual of mental disorders, Fifth Edition.. Am Psychiatric Assoc 2013.

4. Fombonne E. Epidemiology of pervasive developmental disorders. Pediatr Res 2009; 65(6):591–598.

5. Tager-Flusberg H, Paul R, Lord C. Language and Communication in Autism. 2005.

6. Gernsbacher MA, Morson EM, Grace EJ. Language and Speech in Autism. Annu Rev Linguist 2016; 2:413–425.

7. Karolis VR, Corbetta M, Thiebaut de Schotten M. The architecture of functional lateralisation and its relationship to callosal connectivity in the human brain. Nat Commun 2019; 10(1):1417.

8. Frith C, Friston K, Liddle P, Frackowiak R. A PET study of word finding. Neuropsychologia 1991; 29(12):1137–1148.

9. Knecht S, Deppe M, Drager B, Bobe L, Lohmann H, Ringelstein E et al. Language lateralization in healthy right-handers. Brain 2000; 123 (Pt 1): 74–81.

10. Boccadoro S, Cracco E, Hudson AR, Bardi L, Nijhof AD, Wiersema JR et al. Defining the neural correlates of spontaneous theory of mind (ToM): An fMRI multi-study investigation. Neuroimage 2019; 203:116193.

11. Jouravlev O, Kell AJE, Mineroff Z, Haskins AJ, Ayyash D, Kanwisher N et al. Reduced Language Lateralization in Autism and the Broader Autism Phenotype as Assessed with Robust Individual-Subjects Analyses. Autism Res 2020; 13(10):1746–1761.

12. Nielsen JA, Zielinski BA, Fletcher PT, Alexander AL, Lange N, Bigler ED et al. Abnormal lateralization of functional connectivity between language and default mode regions in autism. Mol Autism 2014; 5(1):8.

13. Postema MC, van Rooij D, Anagnostou E, Arango C, Auzias G, Behrmann M et al. Altered structural brain asymmetry in autism spectrum disorder in a study of 54 datasets. Nat Commun 2019; 10(1):4958.

14. Floris DL, Lai MC, Auer T, Lombardo MV, Ecker C, Chakrabarti B et al. Atypically rightward cerebral asymmetry in male adults with autism stratifies individuals with and without language delay. Hum Brain Mapp 2016; 37(1):230–253.

15. Dougherty CC, Evans DW, Katuwal GJ, Michael AM. Asymmetry of fusiform structure in autism spectrum disorder: trajectory and association with symptom severity. Mol Autism 2016; 7:28.

16. van Rooij D, Anagnostou E, Arango C, Auzias G, Behrmann M, Busatto GF et al. Cortical and Subcortical Brain Morphometry Differences Between Patients With Autism Spectrum Disorder and Healthy Individuals Across the Lifespan: Results From the ENIGMA ASD Working Group. Am J Psychiatry 2018; 175(4):359–369.

17. Thompson PM, Jahanshad N, Ching CRK, Salminen LE, Thomopoulos SI, Bright J et al. ENIGMA and global neuroscience: A decade of large-scale studies of the brain in health and disease across more than 40 countries. Transl Psychiatry 2020; 10(1):100.

18. He Y, Chen ZJ, Evans AC. Small-world anatomical networks in the human brain revealed by cortical thickness from MRI. Cereb Cortex 2007; 17(10):2407–2419.

19. Alexander-Bloch A, Giedd JN, Bullmore E. Imaging structural co-variance between human brain regions. Nat Rev Neurosci 2013; 14(5):322–336.

20. Wee CY, Yap PT, Shen D, Alzheimer’s Disease Neuroimaging I. Prediction of Alzheimer’s disease and mild cognitive impairment using cortical morphological patterns. Hum Brain Mapp 2013; 34(12):3411–3425.

21. Yun JY, Boedhoe PSW, Vriend C, Jahanshad N, Abe Y, Ameis SH et al. Brain structural covariance networks in obsessive-compulsive disorder: a graph analysis from the ENIGMA Consortium. Brain 2020; 143(2):684–700.

22. Lerch JP, Worsley K, Shaw WP, Greenstein DK, Lenroot RK, Giedd J et al. Mapping anatomical correlations across cerebral cortex (MACACC) using cortical thickness from MRI. Neuroimage 2006; 31(3):993–1003.

23. Parent A, Carpenter M. Baltimore: Williams and Wilkins; 1995. Human neuroanatomy.

24. Narr KL, Bilder RM, Toga AW, Woods RP, Rex DE, Szeszko PR et al. Mapping cortical thickness and gray matter concentration in first episode schizophrenia. Cereb Cortex 2005; 15(6):708–719.

25. Worsley KJ, Chen JI, Lerch J, Evans AC. Comparing functional connectivity via thresholding correlations and singular value decomposition. Philos Trans R Soc Lond B Biol Sci 2005; 360(1457):913–920.

26. Katz LC, Shatz CJ. Synaptic activity and the construction of cortical circuits. Science 1996; 274(5290):1133–1138.

27. Bi G, Poo M. Distributed synaptic modification in neural networks induced by patterned stimulation. Nature 1999; 401(6755):792–796.

28. Gong G, He Y, Chen ZJ, Evans AC. Convergence and divergence of thickness correlations with diffusion connections across the human cerebral cortex. Neuroimage 2012; 59(2):1239–1248.

29. Catani M, ffytche DH. The rises and falls of disconnection syndromes. Brain 2005; 128(Pt 10): 2224–2239.

30. Menon V. Large-scale brain networks and psychopathology: a unifying triple network model. Trends Cogn Sci 2011; 15(10):483–506.

31. Sha Z, Wager TD, Mechelli A, He Y. Common Dysfunction of Large-Scale Neurocognitive Networks Across Psychiatric Disorders. Biol Psychiatry 2019; 85(5):379–388.

32. Sharda M, Khundrakpam BS, Evans AC, Singh NC. Disruption of structural covariance networks for language in autism is modulated by verbal ability. Brain Struct Funct 2016; 221(2):1017–1032.

33. Sharda M, Foster NEV, Tryfon A, Doyle-Thomas KAR, Ouimet T, Anagnostou E et al. Language Ability Predicts Cortical Structure and Covariance in Boys with Autism Spectrum Disorder. Cereb Cortex 2017; 27(3):1849–1862.

34. Valk SL, Bernhardt BC, Bockler A, Trautwein FM, Kanske P, Singer T. Socio-Cognitive Phenotypes Differentially Modulate Large-Scale Structural Covariance Networks. Cereb Cortex 2017; 27(2):1358–1368.

35. Valk SL, Di Martino A, Milham MP, Bernhardt BC. Multicenter mapping of structural network alterations in autism. Hum Brain Mapp 2015; 36(6):2364–2373.

36. Chow ML, Pramparo T, Winn ME, Barnes CC, Li HR, Weiss L et al. Age-dependent brain gene expression and copy number anomalies in autism suggest distinct pathological processes at young versus mature ages. PLoS Genet 2012; 8(3):e1002592.

37. Writing Committee for the Attention-Deficit/Hyperactivity Disorder, Autism Spectrum Disorder, Bipolar Disorder, Major Depressive Disorder, Obsessive-Compulsive Disorder, and Schizophrenia ENIGMA Working Groups et al. Virtual Histology of Cortical Thickness and Shared Neurobiology in 6 Psychiatric Disorders. JAMA Psychiatry 2021; 78(1):47–63.

38. Yun JY, Kim SN, Lee TY, Chon MW, Kwon JS. Individualized covariance profile of cortical morphology for auditory hallucinations in first-episode psychosis. Hum Brain Mapp 2016; 37(3):1051–1065.

39. Lord C, Risi S, Lambrecht L, Cook EH, Jr., Leventhal BL, DiLavore PC et al. The autism diagnostic observation schedule-generic: a standard measure of social and communication deficits associated with the spectrum of autism. J Autism Dev Disord 2000; 30(3):205–223.

40. Fischl B. FreeSurfer. Neuroimage 2012; 62(2):774–781.

41. Desikan RS, Segonne F, Fischl B, Quinn BT, Dickerson BC, Blacker D et al. An automated labeling system for subdividing the human cerebral cortex on MRI scans into gyral based regions of interest. Neuroimage 2006; 31(3):968–980.

42. Achard S, Bullmore E. Efficiency and cost of economical brain functional networks. PLoS Comput Biol 2007; 3(2):e17.

43. Rubinov M, Sporns O. Complex network measures of brain connectivity: uses and interpretations. Neuroimage 2010; 52(3):1059–1069.

44. Wang J, Wang X, Xia M, Liao X, Evans A, He Y. GRETNA: a graph theoretical network analysis toolbox for imaging connectomics. Front Hum Neurosci 2015; 9:386.

45. Watts DJ, Strogatz SH. Collective dynamics of ‘small-world’ networks. Nature 1998; 393(6684):440–442.

46. Bassett DS, Bullmore E. Small-world brain networks. Neuroscientist 2006; 12(6):512–523.

47. Maslov S, Sneppen K. Specificity and stability in topology of protein networks. Science 2002; 296(5569):910–913.

48. Latora V, Marchiori M. Efficient behavior of small-world networks. Phys Rev Lett 2001; 87(19):198701.

49. Latora V, Marchiori M. Economic small-world behavior in weighted networks. The European Physical Journal B-Condensed Matter and Complex Systems 2003; 32(2):249–263.

50. Nielsen JA, Zielinski BA, Ferguson MA, Lainhart JE, Anderson JS. An evaluation of the left-brain vs. right-brain hypothesis with resting state functional connectivity magnetic resonance imaging. PLoS One 2013; 8(8):e71275.

51. Cohen J. Statistical power analysis for the behavioral sciences. Academic press 2013.

52. Yarkoni T, Poldrack RA, Nichols TE, Van Essen DC, Wager TD. Large-scale automated synthesis of human functional neuroimaging data. Nat Methods 2011; 8(8):665–670.

53. Demetriou EA, DeMayo MM, Guastella AJ. Executive Function in Autism Spectrum Disorder: History, Theoretical Models, Empirical Findings, and Potential as an Endophenotype. Front Psychiatry 2019; 10:753.

54. Demetriou EA, Lampit A, Quintana DS, Naismith SL, Song YJC, Pye JE et al. Autism spectrum disorders: a meta-analysis of executive function. Mol Psychiatry 2018; 23(5):1198–1204.

55. Habib A, Harris L, Pollick F, Melville C. A meta-analysis of working memory in individuals with autism spectrum disorders. PLoS One 2019; 14(4):e0216198.

56. Floris DL, Wolfers T, Zabihi M, Holz NE, Zwiers MP, Charman T et al. Atypical Brain Asymmetry in Autism-A Candidate for Clinically Meaningful Stratification. Biol Psychiatry Cogn Neurosci Neuroimaging 2020.

57. Li T, Hoogman M, Mota NR, Buitelaar J, Vasquez AA, Franke B et al. Dissecting the heterogeneous subcortical brain volume of Autism spectrum disorder (ASD) using community detection. bioRxiv 2020.

58. Westlye LT, Walhovd KB, Dale AM, Bjornerud A, Due-Tonnessen P, Engvig A et al. Differentiating maturational and aging-related changes of the cerebral cortex by use of thickness and signal intensity. Neuroimage 2010; 52(1):172–185.

59. la Fougere C, Grant S, Kostikov A, Schirrmacher R, Gravel P, Schipper HM et al. Where in-vivo imaging meets cytoarchitectonics: the relationship between cortical thickness and neuronal density measured with high-resolution [18F]flumazenil-PET. Neuroimage 2011; 56(3):951–960.

60. Amunts K, Mohlberg H, Bludau S, Zilles K. Julich-Brain: A 3D probabilistic atlas of the human brain’s cytoarchitecture. Science 2020; 369(6506):988–992.

61. Wagstyl K, Larocque S, Cucurull G, Lepage C, Cohen JP, Bludau S et al. BigBrain 3D atlas of cortical layers: Cortical and laminar thickness gradients diverge in sensory and motor cortices. PLoS Biol 2020; 18(4):e3000678.

62. Schmitz J, Fraenz C, Schluter C, Friedrich P, Jung RE, Gunturkun O et al. Hemispheric asymmetries in cortical gray matter microstructure identified by neurite orientation dispersion and density imaging. Neuroimage 2019; 189:667–675.

63. Glasser MF, Van Essen DC. Mapping human cortical areas in vivo based on myelin content as revealed by T1-and T2-weighted MRI. J Neurosci 2011; 31(32):11597–11616.

64. Amaral DG, Anderson MP, Ansorge O, Chance S, Hare C, Hof PR et al. Autism BrainNet: A network of postmortem brain banks established to facilitate autism research. Handb Clin Neurol 2018; 150:31–39.

65. Mosconi MW, Sweeney JA. Sensorimotor dysfunctions as primary features of autism spectrum disorders. Science China Life Sciences 2015; 58(10):1016–1023.

66. Proverbio AM, Brignone V, Matarazzo S, Del Zotto M, Zani A. Gender differences in hemispheric asymmetry for face processing. BMC Neurosci 2006; 7:44.

67. Noesselt T, Driver J, Heinze HJ, Dolan R. Asymmetrical activation in the human brain during processing of fearful faces. Curr Biol 2005; 15(5):424–429.

68. Zhen Z, Yang Z, Huang L, Kong XZ, Wang X, Dang X et al. Quantifying interindividual variability and asymmetry of face-selective regions: a probabilistic functional atlas. Neuroimage 2015; 113:13–25.

69. Kanwisher N, McDermott J, Chun MM. The fusiform face area: a module in human extrastriate cortex specialized for face perception. J Neurosci 1997; 17(11):4302–4311.

70. Hadjikhani N, Joseph RM, Snyder J, Chabris CF, Clark J, Steele S et al. Activation of the fusiform gyrus when individuals with autism spectrum disorder view faces. Neuroimage 2004; 22(3):1141–1150.

71. Cole MW, Reynolds JR, Power JD, Repovs G, Anticevic A, Braver TS. Multi-task connectivity reveals flexible hubs for adaptive task control. Nat Neurosci 2013; 16(9):1348–1355.

72. Lukito S, Norman L, Carlisi C, Radua J, Hart H, Simonoff E et al. Comparative meta-analyses of brain structural and functional abnormalities during cognitive control in attention-deficit/hyperactivity disorder and autism spectrum disorder. Psychol Med 2020; 50(6):894–919.

73. Cardinale RC, Shih P, Fishman I, Ford LM, Muller RA. Pervasive rightward asymmetry shifts of functional networks in autism spectrum disorder. JAMA Psychiatry 2013; 70(9):975–982.

74. Wei L, Zhong S, Nie S, Gong G. Aberrant development of the asymmetry between hemispheric brain white matter networks in autism spectrum disorder. Eur Neuropsychopharmacol 2018; 28(1):48–62.

75. Jeste SS, Geschwind DH. Disentangling the heterogeneity of autism spectrum disorder through genetic findings. Nat Rev Neurol 2014; 10(2):74–81.

76. Masi A, DeMayo MM, Glozier N, Guastella AJ. An Overview of Autism Spectrum Disorder, Heterogeneity and Treatment Options. Neurosci Bull 2017; 33(2):183–193.

